# Targeting alpha-ketoglutarate starvation within the tumor-immune microenvironment rescues T cell effector function in cancer immunotherapy

**DOI:** 10.64898/2026.10.24.761570

**Authors:** Katarina Benejova, Kristina Travis, Deepti Mathur, No joon Song, Zihai Li, Mitch A. Phelps, Elaine Mardis, Liza Makowski, David P. Carbone, Dwight H. Owen, Martin Benej

**Affiliations:** Department of Molecular Medicine and Therapeutics, The Ohio State University Comprehensive Cancer Center - James Cancer Hospital & Solove Research Institute, Columbus, Ohio, United States; Pelotonia Institute for Immuno-Oncology, The Ohio State University Comprehensive Cancer Center - James Cancer Hospital & Solove Research Institute, Columbus, Ohio, United States; Division of Pharmaceutics and Pharmacology, College of Pharmacy, The Ohio State University, Columbus, Ohio, United States; The Steve and Cindy Rasmussen Institute for Genomic Medicine, The Abigail Wexner Research Institute, Nationwide Children’s Hospital, Columbus, Ohio, United States; Department of Pediatrics, The Ohio State University College of Medicine, Columbus, Ohio, United States; Division of Medical Oncology, The Ohio State University Comprehensive Cancer Center - James Cancer Hospital & Solove Research Institute, Columbus, Ohio, United States; The Ohio State University Enterprise for Research Innovation and Knowledge Institute on Aging, Columbus, Ohio, United States

## Abstract

Given its status as the leading cause of global cancer mortality, there is a critical need to enhance immunotherapy strategies for lung cancer to overcome widespread resistance to current treatments. Immunotherapy recruits the patient’s immune system to attack and destroy cancer cells. While highly effective in a subset of patients, chronic exposure of immune cells to cancer cells leads to a progressive loss of effector T cell function and progression of T cell exhaustion; now considered two well-established hallmarks of immunotherapy resistance. The tumor-immune microenvironment (TIME)-driven differentiation into the exhausted T cell phenotype is marked by a loss of metabolic plasticity and epigenome-wide changes in distribution of repressive histone marks such as histone H3 subunit methylation at lysine 27 (H3K27me^3^). Here, we studied how these processes are connected to metabolite availability within the TIME. *In vitro*, chronic antigen stimulation and hypoxia led to rapid loss of effector function, associated with a suppression of alpha-ketoglutarate (aKG) intracellular concentrations in T cells. Supplementation of cell-permeable aKG led to a rescue of effector cytokine production and to improved cytotoxic capacity despite chronic antigen stimulation and hypoxia. Increased intracellular aKG levels restored the activity of H3K27 demethylases UTX and JMJD3, consistent with epigenome-wide decreased H3K27me^3^. *In vivo*, dietary supplementation with commercially available calcium-aKG (Ca-aKG) supplements led to an elevation of serum and intratumoral aKG levels associated with a significant increase of immunotherapy effectiveness in mouse models of NSCLC. Clinical samples from patients with NSCLC contained significantly decreased serum aKG levels in older NSCLC patients (>65 years of age). Analysis identified a novel association between serum aKG levels and poor immunotherapy response. Thus, our work demonstrates that intratumoral metabolite “starvation” can be therapeutically exploited to improve immunotherapy outcomes in non-small cell lung cancer (NSCLC).

## INTRODUCTION

Non-small cell lung cancer (NSCLC) is the leading cause of cancer-related death worldwide. Immune checkpoint inhibitors (ICIs) have transformed outcomes for a small subset of patients with NSCLC (1–3). Unfortunately, fewer than 20% of patients with advanced NSCLC without an actionable genetic alteration (i.e. *EGFR, ALK, ROS1*) will survive for 5 years after diagnosis despite receiving ICI treatment (4). These poor outcomes highlight a critical unmet need to understand why ICIs fail to provide durable benefit and identify strategies to improve patient survival.

Effector CD8^+^ T cells (Teff) are an integral part of the anti-tumor immune response. However, tumor-immune microenvironment (TIME) factors such as chronic antigen exposure, hypoxia, and inhibitory signals drive their differentiation into a dysfunctional, exhausted state (5). Exhausted T cells (Tex) are functionally, transcriptionally, and epigenetically distinct from the Teff cells and are characterized by a progressive loss of effector function and proliferation, sustained upregulation and co-expression of inhibitory receptors, and by the inability to transition to the quiescent antigen-independent memory T cell state (6, 7). While initially considered to be a uniform population of dysfunctional T cells, it is now accepted that Tex are a heterogenous population with distinct stages of dysfunction characterized by unique transcription and epigenetic patterns (8). In early phases of chronic antigen exposure, Teff gradually lose their effector function and differentiate into progenitor Tex cells, a unique T cell population exhibiting stem-like properties including self-renewal, proliferation and differentiation into Tex (9, 10). The progenitor Tex display high levels of T cell factor 1 (TCF-1), SLAM Family Member 6 (SLAMF6), and C-X-C chemokine receptor type 5 (CXCR5) receptor expression; and a gradual loss of interleukin-2 (IL-2) and tumor necrosis factor-α (TNFα) production, consistent with compromised cytotoxicity at this early stage of ehxaustion (11). Of note, progenitor Tex are the target population for reinvigoration by anti-PD-1 immune checkpoint inhibitors (ICIs) (12). Continued antigen exposure leads to further transcriptome- and effector function-level changes most prominent of which is the loss of TCF-1 expression and interferon γ (IFNγ) production, leading to differentiation into the terminally exhausted Tex, which are resistant to ICIs (13, 14). Thus, Tex cells pose a barrier to effective cancer immunotherapy and therefore TIME factors driving Teff-to-Tex differentiation are under intense investigation.

Loss of metabolic plasticity has been shown to be directly linked to T cell exhaustion (15–17). Following antigen stimulation, naïve CD8^+^ T cells undergo massive proliferation and clonal expansion to differentiate into Teff cells (18). To fulfill the associated increased energetic demands and to provide biosynthetic intermediates required to support elevated protein and nucleotide synthesis, activated CD8^+^ T cells reprogram their metabolism from oxidative phosphorylation (OXPHOS) to aerobic glycolysis, orchestrated by the phosphatidylinositol-3-kinase–protein kinase B–mammalian target of rapamycin (PI3K–Akt–mTOR) pathway (19, 20). In contrast, effector function- and proliferation-deficient Tex cells exhibit suppressed mitochondrial OXPHOS and aerobic glycolysis pathways, associated with metabolic deficiency and altered mitochondrial mass and membrane potential (21–23). These findings signal that metabolite availability and/or deprivation have a critical impact on T cell homeostasis and functionality, and imply that metabolic reprogramming by metabolite supplementation can be a promising therapeutic avenue to restore T cell function and improve immunotherapy outcomes (24).

Here, we demonstrate that TIME suppresses the intracellular levels of alpha-ketoglutarate (aKG), a key TCA cycle metabolite that we and others have found to be essential for maintaining Teff function (25, 26). Under normoxic conditions, aKG is produced by the classical TCA reaction of oxidative decarboxylation of isocitrate by isocitrate dehydrogenases (IDH), or alternatively from glutamine through the mitochondrial enzymes glutaminase (GLS) and glutamate dehydrogenase (GLUD) (27). Within the hypoxic TIME however, hypoxia-inducible factor 1 (HIF-1)-mediated induction of pyruvate dehydrogenase kinases (PDHKs) decreases pyruvate dehydrogenase flux and carbon entry into TCA reactions, thus leading to suppression of aKG production (28). aKG is an obligatory substrate for cellular dioxygenases such as Jumonji C domain (JmjC)- containing lysine demethylases KDM 2–7, HIF prolyl hydroxylases (PHDs), and ten-eleven translocation hydroxylases (TET1–3) (29). Tex cells display enriched levels of the repressive histone mark histone H3 subunit methylation at lysine 27 (H3K27me^3^) in loci associated with the Teff function (30, 31). H3K27me^3^ is exclusively removed by aKG-dependent histone demethylases *kdm6a/b* (UTX/JMJD3), providing a functional link between TIME-depleted aKG levels and reduced Teff function.

To evaluate the rationale to therapeutically elevate aKG levels to improve ICI outcomes, we used non-small cell lung cancer (NSCLC) models. Here, we investigated the role of aKG levels in restoring the cytotoxic function of infiltrating Teffs. Interestingly, chronically stimulated CD8^+^ T cells exhibited intracellular aKG deficiency even in normoxia. Using a multi-omic approach, we identified that supplementation with calcium alpha-ketoglutarate (Ca-aKG), a highly bioavailable salt form of aKG sustains an active Teff transcription program and cytotoxic function, and removes the genome-wide H3K27me^3^ epigenetic signature in chronically stimulated hypoxic CD8^+^ T cells. The use of aKG supplementation in animal models and humans has been studied for many healthcare applications including aging, osteoporosis and liver disease (32–40). However, these studies found that potential clinical applications of supplementation with standard aKG formulation are limited due to its weakly acidic pH which restricts its permeability through cell membranes; and due to its very short half-life (typically less than 5 minutes) (41–43). Upon administration, more than 80% of soluble aKG is rapidly removed from the bloodstream, and the remaining aKG is absorbed by first-pass metabolism in enterocytes and liver, and converted to amino acids such as glutamine, proline or leucine (44, 45). Esterified cell-permeable variants such as dimethyl alpha-ketoglutarate (Dm-aKG) are commonly used for *in vitro* studies, but these variants are associated with high levels of metabolic toxicity; their hydrolysis is associated with release of methanol and formaldehyde, which prevent their adaptation for therapeutic use (46). In summary, these findings demonstrate that although clinically sound, therapeutic elevation of intratumoral aKG through dietary supplementation is challenging.

In contrast, Ca-aKG is a salt formulation of aKG in which two protons were removed to facilitate calcium binding, leading to improved stability and bioavailability, and allowing longer retention in the body (47, 48). Dietary Ca-aKG supplements are commercially available without prescription and have long been used in healthcare to treat a variety of conditions including osteopenia without significant toxicities (36, 49, 50), suggesting that translation to anti-tumor therapy could be feasible. We observed that NSCLC tumor-bearing mice (inoculated with lung adenocarcinoma-derived CMT167 or squamous cell carcinoma-derived KLN 205) fed with 4% Ca-aKG diet exhibited increased serum and intratumoral aKG levels. Importantly, Ca-aKG augmented ICI effectiveness without significant observed adverse effects.

Notably, NSCLC disproportionately affects older individuals; the median age at diagnosis is 71 years and more than three quarters of all new patients are older than 65 years (51, 52). We identified that serum aKG levels were significantly lower in older NSCLC patients compared to younger adults with NSCLC. Critically, reduced serum aKG levels correlated with poor ICI outcome. Based on our results, we posit that combining aKG dietary supplementation with ICI therapy could improve patient outcomes for those suffering with NSCLC. To identify patient populations most likely to benefit from dietary aKG supplementation-based augmentation of their ICI therapy, we developed an *in silico* model parametrized on pre-clinical and clinical data that predicts NSCLC patients without an actionable genetic alteration and with serum aKG levels below 8 μM will achieve the maximum ICI benefit of Ca-aKG supplementation. Dietary Ca-aKG supplements thus may potentially increase ICI outcomes not only for NSCLC but other solid cancers that are ICI resistant.

## RESULTS

### Chronic antigen exposure and hypoxia promote T cell exhaustion by decreasing cellular aKG levels

To evaluate the effects of hypoxic TIME on the T cell metabolism, we assayed TCA metabolite utilization in anti-CD3/CD28-activated murine CD8^+^ T cells under chronic antigen exposure (10 days; T cell receptor (TcR) re-stimulated every two days with 3 μg of anti-CD3 antibody; adopted from (53)) and hypoxia (1% O_2_) (**Fig. 1A**). Compared to normoxic controls (21% O_2_), hypoxic T cells exhibited decreased utilization of multiple TCA metabolites, most prominent of which was aKG (**Fig. 1B, Supp. Fig. S1A**). We confirmed that the observed changes of metabolic preference under chronic TcR stimulation and hypoxia are associated with a reduction of effector cytokine TNFα and IFNγ secretion (**Fig. 1C-D**). To determine whether mitochondrial dysfunction associated with chronic TcR stimulation decreases the aKG availability, we assayed intracellular aKG levels in activated CD8^+^ T cells under acute (TcR stimulated once) or chronic (TcR re-stimulated every two days for 10 days) antigen exposure. Of note, chronic TcR stimulation was associated with a significant reduction of intracellular aKG levels (p-value=0.008296 by Student’s t-test) (**Fig. 1E**).

**Fig. 1.**
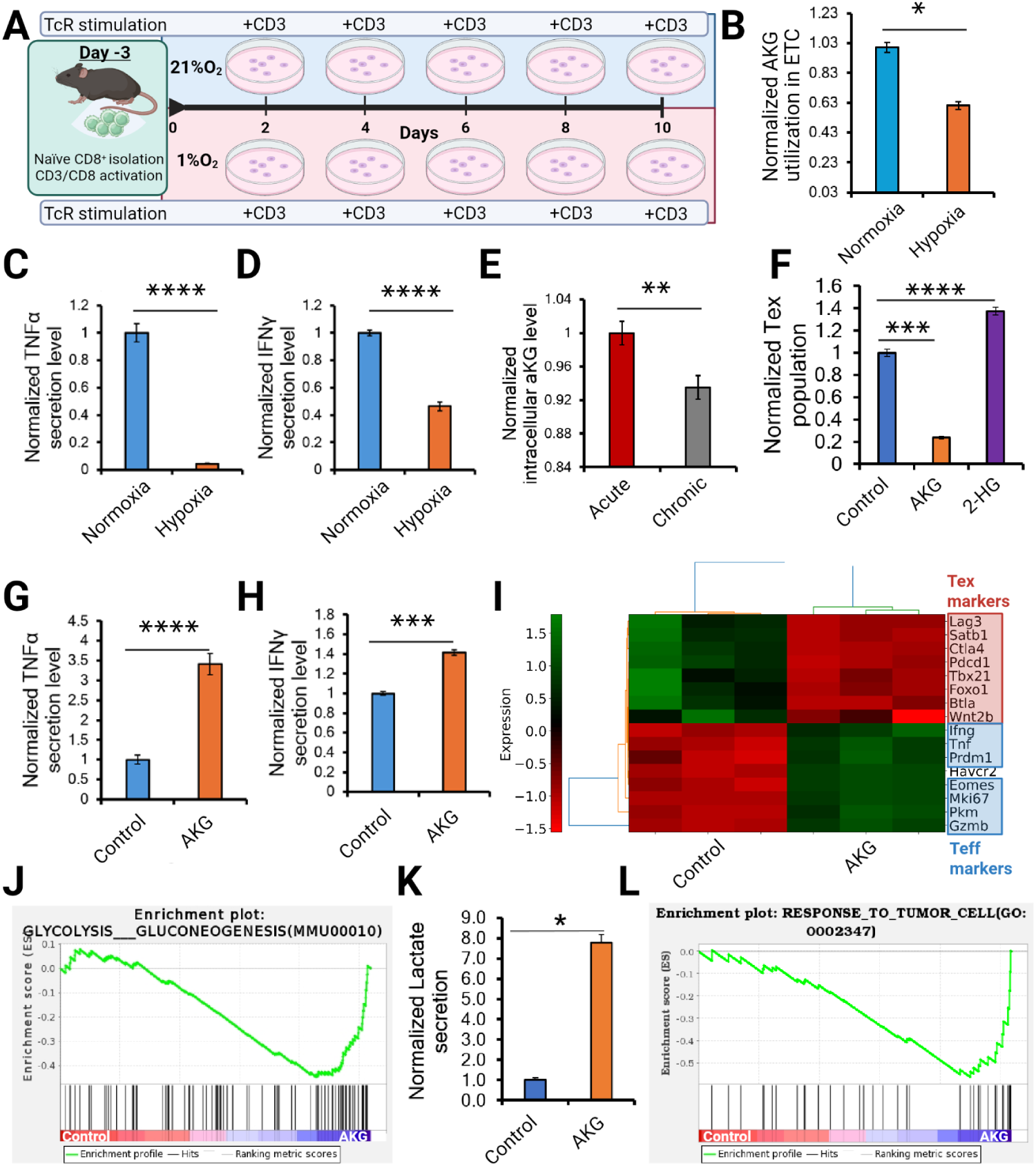
Elevation of intracellular aKG rescues Teff function. **(A)** Schematic overview of *in vitro* T cell exhaustion assay, adapted from (53). Naïve murine CD8^+^ T cells are activated using anti-CD3/CD28 antibody cocktail for 72 hours, followed by culturing under chronic TcR stimulation (3 μg/ml anti-CD3 antibody restimulated every two days) for 10 days in normoxia (21% O2) or hypoxia (1% O2). **(B)** aKG utilization in hypoxic CD8^+^ T cells under chronic TcR stimulation normalized to the normoxic control, as determined by Mitoplate S-1 assay (n=3 biological replicates). **(C-D)** Quantification of effector cytokine TNFα (C) and IFNγ (D) secretion from CD8^+^ T cells under chronic TcR stimulation in normoxia or hypoxia for 10 days by ELISA (n=3). **(E)** Intracellular aKG levels in normoxic CD8^+^ T cells stimulated once (*Acute*) or every two days (*Chronic*) for 10 days (n=3). **(F)** Normalized exhausted T cell (Tex) populations (PD-1^+^/Tim3^+^/Lag3^+^) after 3 days of hypoxia and chronic TcR stimulation ± cell-permeable aKG (dimethyl-aKG) (*purple*) or its competitive inhibitor 2-L-hydroxyglutarate (2-HG, *orange*), (both at 10 mM); determined by multispectral flow cytometry (n=5 biological replicates). **(G-H)** Quantification of effector cytokine TNFα (G) and IFNγ (H) secretion in CD8^+^ T cells under chronic TcR stimulation and hypoxia for 10 days ± 10 mM cell-permeable aKG by ELISA (n=3). **(I)** RNA-seq analysis of CD8^+^ T cells ± 10 mM cell-permeable aKG; effector T cell (Teff) / Tex-specific genes are shown. **(J)** GSEA analysis calculated for Control vs cell-permeable aKG (AKG) groups revealed significant upregulation of the glycolysis/gluconeogenesis pathway (KEGG: MU00010). **(K)** Extracellular lactate levels in normoxic CD8^+^ T cells after 10 days of hypoxia and chronic TcR stimulation ± cell-permeable aKG, as determined by commercial colorimetric assay (n=3). **(L)** GSEA analysis calculated for Control vs cell-permeable aKG (AKG) groups revealed significant upregulation of the “Response to tumor” pathway (GO:002347). P values were calculated by Student’s t-test. *P<0.05, **P<0.01, *** P < 0.001, **** P < 0.0001.

To determine whether elevation of the intracellular aKG levels could rescue the Teff function, we supplemented activated CD8^+^ T cells under chronic antigen stimulation and hypoxia with dimethyl alpha-ketoglutarate (Dm-aKG), a cell-permeable ester of aKG (54). We found that intracellular aKG pool elevation is associated with a significant decrease of the Tex population (p-value=0.0007 by Student’s t-test), while treatment with L-2-hydroxyglutarate (2-HG), a competitive aKG inhibitor, had the opposite effect to boost Tex cell numbers (**Fig. 1F**). We next assayed the effector function and identified a significant elevation of effector cytokine TNFα and IFNγ secretion (p-value=0.000065 and 0.007497, respectively, by Student’s t-test) (**Fig. 1G-H**), consistent with a rescue of cytotoxic capacity (**Supp. Fig. S1B)** in hypoxic CD8^+^ T cells under chronic TcR stimulation. On the transcriptome level, while chronic antigen stimulation led to suppression of proliferation- and effector function-associated genes; Dm-aKG-treated T cells displayed an active Teff-specific expression pattern and a suppressed Tex-specific co-inhibitory marker expression pattern (**Fig. 1I**). The elevated effector function resulted in an increased reliance on the glycolytic metabolism as evidenced by our Gene Set Enrichment Analysis (GSEA) (**Fig. 1J**), and validated by increased extracellular lactate levels (**Fig. 1K**). Additional pathways related to response to nutrients, response to tumor cells, as well as T cell activation and proliferation pathways were enriched in response to aKG pool elevation (**Fig. 1L**). Taken together, these findings indicate that TIME-mediated repression of T cell function can be reversed in an aKG-dependent manner.

### Elevation of intracellular aKG rescues Teff function by affecting the epigenome

We next sought to investigate the mechanism of aKG-mediated rescue of Teff function. Chronic TcR stimulation has been shown to repress mitochondrial oxidative metabolism through PPAR-gamma coactivator 1α (PGC-1α) suppression, leading to a persistent loss of mitochondrial function and mass (23). To determine whether aKG supplementation affects mitochondrial mass, we assayed voltage-dependent anion channel (VDAC) levels in Dm-aKG-treated T cells under chronic TcR stimulation (**Supp. Fig. S1C**). No significant difference in mitochondrial mass was observed suggesting that aKG supplementation does not rescue the Teff function via mitochondrial mass elevation. Consistent with these findings, no changes in PGC-1α expression were observed between the groups. However, aKG plays a vital role in multiple cellular processes that may directly affect the metabolic plasticity, including redox balance and epigenetic modification (55, 56). Chronic TcR stimulation has been shown to promote reactive oxygen species (ROS) accumulation in the mitochondria of CD8^+^ T cells, leading to increased levels of lipid peroxidation and uncoupling of oxidative phosphorylation (53). Moreover, hypoxic TIME is associated with increased superoxide levels, triggered by disruption of mitochondrial electron transport chain (57, 58). aKG displays antioxidative capacity by upregulation of endogenous antioxidant enzymes such as superoxide dismutase (SOD1-2), catalase (CAT), and suppression of hydrogen peroxide (H_2_O_2_) levels and by serving as glutamate source to support glutathione synthesis (59, 60). This led to the expectation that at least a part of the effects of aKG elevation on Teff function might be mediated through ROS scavenging. However, on the transcriptome level, Dm-aKG-treated T cells did not display biologically relevant upregulation of SOD1, SOD2, or CAT expression (0.4-fold, 0.11-fold and 0.4-fold, respectively). To further discern whether aKG rescues the Teff function by decreasing intracellular ROS levels, we assayed the levels of superoxide formation in CD8^+^ T cells under chronic TcR stimulation. At the Dm-aKG concentration evaluated throughout this study, elevated aKG levels did not significantly affect superoxide levels in normoxia, or hypoxia (**Supp. Fig. S1D**). These findings imply that aKG elevation does not restore Teff function by direct effects on the mitochondrial mass or redox homeostasis.

The progressive nature of TIME-mediated differentiation of effector T cells into terminally exhausted T cells is regulated by a major epigenomic overhaul (61–63). In addition to decreased chromatin accessibility, multiple studies have reported enriched levels of the repressive histone mark H3K27me^3^ in loci associated with the effector function and proliferation, and a permissive epigenetic signature in loci encoding terminal exhaustion-specific co-inhibitory receptors, such as *Tox* and *Lag3* (30, 31, 63, 64). aKG is an obligatory substrate for the activity of major H3K27 histone demethylases UTX and JMJD3 (encoded by *kdm6a-b*) (**Fig. 2A**) (65). To determine whether the activity of UTX and JMJD3 affects T cell differentiation into the terminally exhausted phenotype, we assayed the progressive loss of TCF-1 expression in Tox^+^ CD8^+^ T cells under persistent TcR stimulation and hypoxia treated with UTX/JMJD3 inhibitor GSK-J4. Of note, we observed a significant elevation of the terminally exhausted Tox^+^ TCF-1^-^ T cell population in response to UTX/JMJD3 inhibition (p=0.001254 by Student’s t-test) (**Fig. 2B**). This implies that elevation of aKG levels could reverse one of the causes of T cell dysfunction by reactivating the UTX/JMJD3 demethylases, which would support an active effector T cell epigenetic signature. We have therefore assayed the enzymatic activity of these two enzymes in activated CD8^+^ T cells under chronic antigen stimulation and hypoxia in the presence or absence of Dm-aKG. We confirmed that Dm-aKG supplementation elevated the enzymatic activity of UTX/JMJD3 demethylases in these T cell exhaustion-promoting conditions (**Fig. 2C**). To validate that revived JMJD3/UTX activity leads to a reduction of H3K27me^3^ levels in activated CD8^+^ T cells under chronic antigen exposure and hypoxia, we used a single-cell western (scWestern) approach. We observed an almost complete elimination of the H3K27me^3^ abundance in Dm-aKG treated CD8^+^ T cells (**Fig. 2D**), supporting the model that elevation of aKG levels may affect the epigenetic processes that regulate TIME-mediated Tex differentiation.

**Fig. 2.**
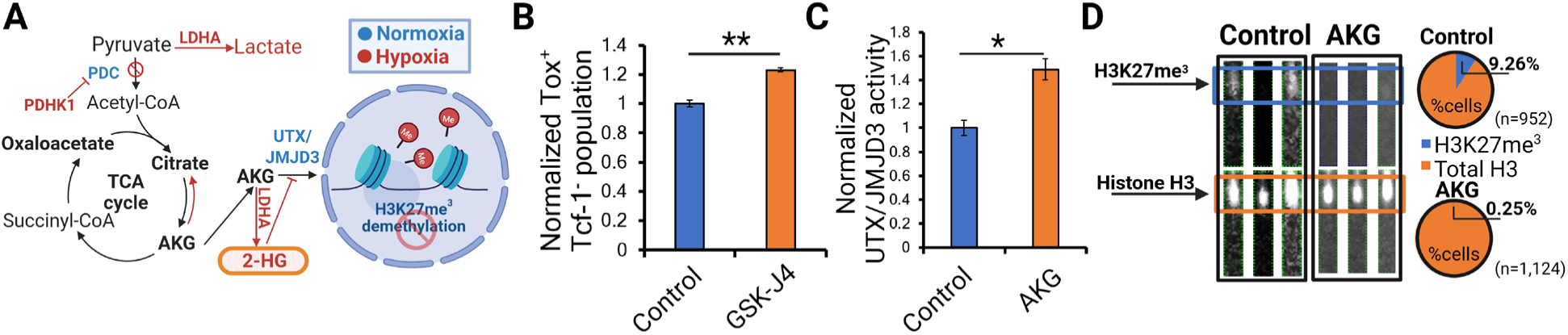
Elevation of intracellular aKG rescues Teff function by affecting the epigenome. **(A)** Hypoxic rewiring of TCA metabolism produces less aKG, while also promoting lactate dehydrogenase A (LDHA)-catalyzed conversion of aKG to its competitive inhibitor 2-L-hydroxyglutarate (2-HG). 2-HG directly inhibits the UTX/JMJD3 demethylases which remove H3K27me^3^. **(B)** Flow cytometry-based quantification of terminally exhausted T cell population (Tox^+^/TCF-1^-^) from CD8^+^ T cells treated with chronic antigen and hypoxia for 10 days ± 10 μM GSK-J4; a UTX/JMJD3 inhibitor added on Day 6. (n=2-4). **(C)** Quantification of UTX/JMJD3 demethylase activity by a commercial assay. Nuclear extracts from CD8^+^ T cells treated with chronic antigen and hypoxia for 10 days were assayed at 2.5 μg each in triplicates and the ability of UTX/JMJD3 to remove plate-bound H3K27me^3^ was assessed by a colorimetric antibody-based assay. **(D)** Representative lanes from scWestern analysis of CD8^+^ T cells under chronic antigen exposure and hypoxia ± 10 mM Ca-aKG for 10 days. T cells were assayed on an scWestern chip and on-chip chromatin digestion was performed to make H3K27me^3^-modified histones physically accessible for scWestern analysis. Each lane shows H3K27me^3^ level (*blue*) and the respective histone H3 level (*orange*) for the same cell. On the right, pie charts show % of H3K27me^3^ positive T cells (*blue*) normalized to total histone H3 levels (*orange*) in the Control group (*top*; n=952 CD8^+^ T cells analyzed) vs Ca-aKG-supplemented group (*bottom*; n=1,124 CD8^+^ T cells analyzed). P values were calculated by Student’s t-test. *P<0.05; **P<0.01.

### Supplementation with Ca-aKG improves ICI response in a murine NSCLC tumor model *in vivo*

Our transcriptomic and mechanistic assays indicate that TIME-mediated aKG dysregulation contributes to the formation of Tex-specific epigenomic signature by promoting H3K27me^3^ enrichment. This implies that therapeutic elevation of aKG levels may improve immunotherapy response by affecting the epigenetic processes that regulate TIME-mediated Tex differentiation. However, *in vivo* use of Dm-aKG is limited due to metabolic toxicity (46). Moreover, potential clinical applications of supplementation with regular aKG formulation are limited due to its weakly acidic pH which restricts its permeability through cell membranes; as well as its very short half-life (< 5 minutes) (41–43).

To address this issue, we used calcium aKG (Ca-aKG); a widely available salt formulation of aKG in which two protons were removed to facilitate calcium binding, leading to improved stability and bioavailability, and allowing longer retention in the body (47) (**Fig. 3A**). We have formulated 4% Ca-aKG-enriched mouse diet into a regular Teklad 7912 diet commonly used in animal facilities. This represents a human equivalent dose (66) of 3.4 g/day; which equals approximately 56% of the dose safely used in a human osteopenia clinical trial (45). Because NSCLC disproportionately affects older patients, we used aged wild-type C57BL/6J mice (35-week-old), inoculated with CMT167 cells in the flank. Upon detecting palpable tumors, the mice were randomized into cohorts fed with regular diet or 4% Ca-aKG diet. After 7 days, serum and intratumoral aKG levels were measured by LC-MS, confirming that dietary Ca-aKG supplementation feasibly elevates serum and intratumoral aKG levels in tumor-bearing mice (**Fig. 3B-C**). Interestingly, spatial metabolomics analysis (MALDI-ToF-based imaging) of tumor sections from both diet cohorts revealed that tumor regions enriched for aKG following Ca-aKG diet did not colocalize with elevated levels of glucose, glutamine, or glutathione (**Fig. 3D**). To evaluate whether increased intratumoral aKG levels were associated with improved immunotherapy response, we grew CMT167 tumors in the flanks of C57BL/6J mice. Upon reaching 100 mm^3^ tumor volumes, the mice were randomized into regular diet or 4% Ca-aKG diet cohorts. Three days later, the mice from both cohorts received two cycles of either rat isotype IgG control or anti-PD-1 antibody (70 μg, I.P.); three days apart (**Fig. 3E**). Because the CMT167 tumor model is generally responsive to ICI therapy (67), we used a suboptimal monotherapy dose that alone does not delay tumor growth (70 μg; **Fig. 3F**), allowing a feasible quantification of the therapeutic benefit of the Ca-aKG diet. Interestingly, dietary Ca-aKG supplementation was associated with a significantly increased anti-PD-1 response compared to isotype control (p-value=0.000542 by Student’s t-test) (**Fig. 3F**).

**Fig. 3.**
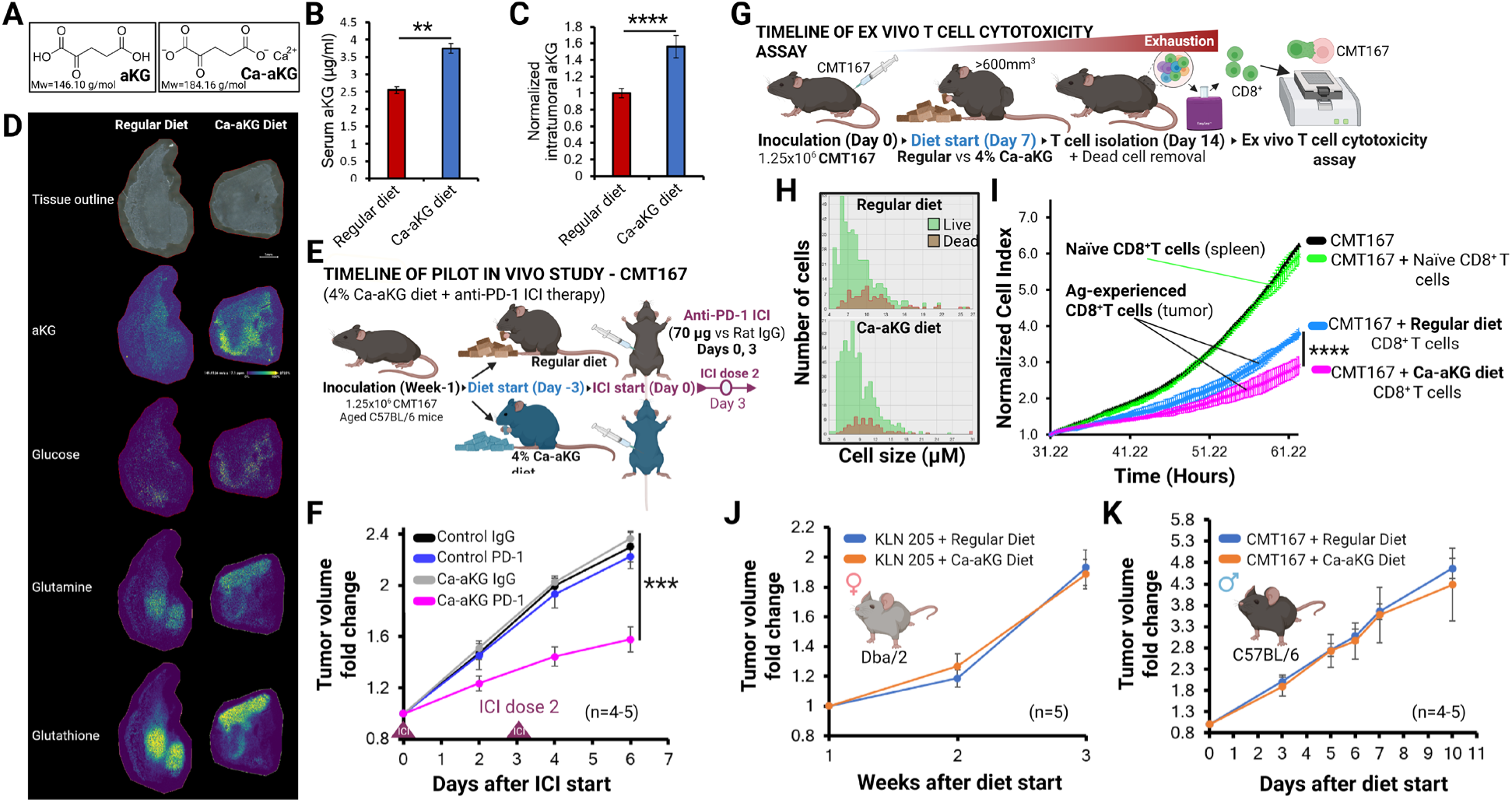
Supplementation with Ca-aKG improves ICI response in a murine NSCLC tumor model. **(A)** Chemical structure of aKG and Ca-aKG. **(B-C)** Serum aKG levels (C) or intratumoral aKG levels (D) in 35-week-old C57BL/6J mice bearing CMT167 tumors fed with regular diet or 4% Ca-aKG cohorts, as quantified by LC-MS (n=4-5). **(D)** Representative spatial metabolomics analysis of aKG, glucose, glutamine and glutathione distribution in regular diet vs Ca-aKG diet tumors, as determined by MALDI-ToF analysis of tumor sections. **(E)** Overview of Ca-aKG-enriched diet regimen. **(F)** CMT167 tumor growth in female C57BL/6J mice (aged 35 weeks) on regular diet or 4% Ca-aKG diet, treated with 70 μg of anti-PD-1 antibody on Day 0 and Day 3 (n=4-5). Error bars are SEM. **(G)** Overview of *ex vivo* T cell cytotoxicity assay. **(H)** Viability of tumor-infiltrating CD8^+^ T cells isolated and pooled from Regular diet or Ca-aKG diet tumors (n=4). **(I)** Quantification of cytotoxic activity of antigen (Ag)-experienced T cells from Regular diet (*blue*) or Ca-aKG diet (*pink*) cohort CMT167 tumors. Naïve CD8^+^ T cells (*green*) were isolated from spleens of tumor-bearing mice as a control. *Black* curve shows CMT167 proliferation without any T cells added. **(J-K)** Daily 4% Ca-aKG diet alone did not affect the growth of: (J) KLN 205 tumors grown in female Dba/2 mice (n=5); or (K) CMT167 tumors grown in male C57BL/6J mice (n=4-5). Control mice received regular diet; all mice are ∼35 weeks old. P values were calculated by Student’s t-test. * P < 0.05; ** P < 0.01; *** P < 0.001; **** P < 0.0001.

Multispectral flow cytometry-based immunophenotyping analysis revealed that the improved ICI response was not associated with increased T cell infiltration into the tumors (**Supp. Fig. S1E**). To validate that Ca-aKG supplementation rescues the anti-PD-1 response by affecting the Teff function, we developed a novel assay measuring the *ex vivo* cytotoxic capacity of antigen-experienced infiltrating CD8^+^ T cells isolated from regular diet or Ca-aKG cohort tumors (**Fig. 3G**). Here, antigen-experienced infiltrating CD8^+^ T cells were harvested from tumors by immunomagnetic separation and upon confirming the viability and dead cell removal, were co-cultured with the cancer cell lines used to grow the tumors at a pre-determined effector-to-target ratio. The cytotoxic capacity was evaluated in real time as a function of cancer cell proliferation using an Agilent xCELLigence RTCA S16 platform (**Fig. 3G**). While CD8^+^ T cells isolated from both cohort tumors showed comparable population size and viability (**Fig. 3H**), the capacity of antigen-experienced CD8^+^ T cells to kill CMT167 cells *ex vivo* was significantly higher in T cells isolated from the Ca-aKG cohort tumors compared to that of the CD8^+^ T cells isolated from the regular diet tumors (p-value=0.00006788 by T-test) (**Fig. 3I**). Similar results were observed using less immunogenic squamous cell carcinoma-derived KLN 205 tumors grown in the Dba/2 mouse strain fed with either regular diet or Ca-aKG enriched diet. While studies show that squamous cell carcinomas are typically associated with inferior ICI outcomes compared to adenocarcinomas (68, 69), we demonstrate that CD8^+^ T cells isolated from Ca-aKG diet KLN 205 tumors display significantly increased cytotoxic activity compared to CD8^+^ T cells isolated from the regular diet tumors (p-value=0.00005577 by Student’s t-test) (**Supp. Fig. S1F**). Consistent with expectations for increased metabolic demand for glycolysis in highly activated Teff cells, our spatial metabolomics analysis revealed elevated levels of lactate in the Ca-aKG tumors compared to the regular diet cohort tumors (**Supp. Fig. S1G**).

Ca-aKG supplementation has not been evaluated in the context of cancer immunotherapy in NSCLC. To explore potential adverse effects, we demonstrated that cell-permeable aKG supplementation does not affect CMT167 or KLN 205 cancer cell proliferation *in vitro* (**Supp. Fig. S1H-I**); and that the 4% Ca-aKG diet alone does not stimulate KLN 205 or CMT167 tumor growth in Dba/2 or C57BL/6J female or male mouse hosts *in vivo* (**Fig. 3J-K**). Moreover, complete necropsy and blood analysis showed no obvious hematological, hepatic, renal, or electrolyte toxicities nor any evidence of histopathologic tissue changes (**Supp. Fig. S1J-L**). Elevation of intratumoral aKG levels may affect other dioxygenases such as HIF prolyl hydroxylases which may affect the metabolite availability within the TIME. We measured HIF-1α levels in Regular diet vs 4% Ca-aKG cohort CMT167 tumors grown in 35-week-old C57BL/6J mice by immunohistochemistry and observed no significant intratumoral HIF-1α level changes (**Supp. Fig. S1M**).

### Decreased serum aKG levels predict poor immunotherapy outcomes in NSCLC

Previous studies in colorectal cancer established that aKG levels are reduced in tumor specimens relative to adjacent normal tissue, providing a link between aKG levels and ICI response in colorectal cancer (25). However, aKG levels in NSCLC progression and ICI response have not been reported. We therefore evaluated whether serum aKG levels are dysregulated in NSCLC patients compared to healthy donors by LC-MS analysis. Interestingly, we observed no significant difference in mean serum aKG levels between the two groups (5,584 ng/ml in pre-treatment NSCLC samples vs 5,386 ng/ml in healthy donors; n=15). Serum aKG levels in healthy individuals were reported to naturally decline with increasing chronological age (70). NSCLC is disproportionately diagnosed in older individuals and the median age at diagnosis is 71 years (51). To discern whether increasing chronological age affects serum aKG levels in NSCLC patients, we analyzed serum aKG levels in patients with NSCLC prior to receiving their ICI treatment in a prospective observational study (NCT06630429); median age at diagnosis was 67 years. We identified that serum aKG levels are significantly lower in older NSCLC patients (> 65 years of age) compared to younger patients (< 65 years) (**Fig. 4A**). To evaluate whether serum aKG levels correlate with ICI response in NSCLC, we conducted a retrospective analysis of a cohort of patients treated with ICIs (pembrolizumab, nivolumab, durvalumab) between 2015-2025 with blood specimens available (TCC Cohort). We identified that low serum aKG levels associated with poor overall survival in NSCLC patients treated with ICIs (**Fig. 4B**). These findings suggest that the majority of NSCLC patients may have already decreased aKG levels at the start of their ICI treatment which provides the rationale to therapeutically elevate aKG levels to improve ICI outcomes in NSCLC. To model whether older patients would be particularly responsive to Ca-aKG supplementation, we supplemented 7-week-old and 35-week-old C57BL/6J mice with 4% Ca-aKG enriched diet for 2 weeks and assayed the endpoint serum aKG levels by LC-MS. We found that while Ca-aKG-enriched diet leads to sustainable elevation of serum aKG in both age cohorts; 35-week-old mice exhibit a higher increase of mean serum aKG levels compared to 7-week-old mice receiving the same Ca-aKG dose (1.9-fold vs 1.48-fold, respectively, **Fig. 4C**).

**Fig. 4.**
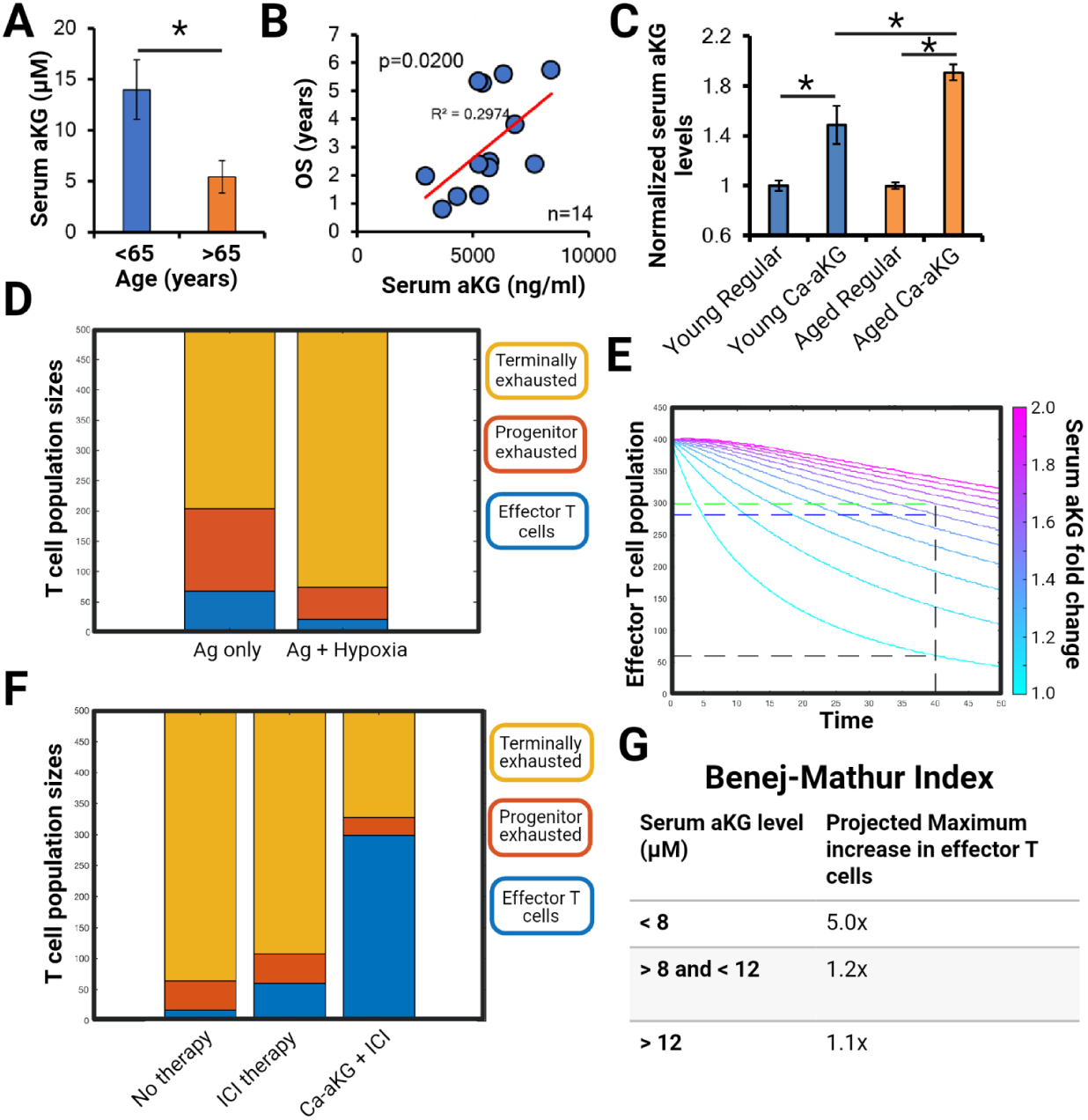
Decreased serum aKG levels predict poor immunotherapy outcomes in NSCLC. **(A)** Serum aKG levels in a prospective cohort of OSU NSCLC patients prior to receiving their ICI treatment (n=12). P values calculated by t test. *P<0.05 **(B)** Serum aKG in a retrospective cohort of patients with NSCLC receiving ICI treatment at OSU between 2015-2025 (n=14). F p-value calculated by F-test for linear regression. **(C)** Endpoint serum aKG levels in 7-week-old (Young) or 35-week-old (Aged) C57BL/6J mice fed with regular diet or Ca-aKG diet for 2 weeks. Ca-aKG cohorts in both age groups were normalized to their respective regular diet controls. P values were calculated by Student’s t-test. *P<0.05. **(D)** Predicted relative proportion of each T cell population at the same point in time under persistent antigen exposure only or under persistent antigen exposure and hypoxia. **(E)** Predicted effector T cell population size changes over time upon ICI and Ca-aKG supplementation. Colors represent relative simulated Ca-aKG levels compared to baseline. Therapy was initiated in a therapeutic window prior to steady state exhaustion (t=0). **(F)** Predicted relative proportion of each T cell population at the same point in time under no therapy, ICI alone, or Ca-aKG combined with ICI. Experimentally validated 1.6-fold elevation of serum aKG in response to dietary Ca-aKG supplementation in mouse models was used to calculate the combination therapy effects. **(G)** Projected increase in effector T cell population due to Ca-aKG supplementation, for each stratification of baseline serum aKG levels. Patients with serum aKG levels < 8μM are predicted to achieve the greatest maximum benefit of Ca-aKG supplementation.

Finally, to provide a better understanding of whether aKG elevation may affect the effector function of infiltrating CD8^+^ T cells in NSCLC, we developed a structured finite-state mathematical model using ordinary differential equations (ODEs) to simulate T cell transitions with respect to the TIME. Building on prior work by our group and others (71, 72), we developed a computational tumor model with spatial irregularities in perfusion and consequent heterogeneity, using Buffa hypoxia scores (73) in NSCLC patients obtained from The Cancer Genome Atlas (TCGA) as surrogate inputs for real-world hypoxia levels in NSCLC (74). Concordant with our experimental data, our model demonstrates that under persistent TcR stimulation, hypoxia accelerates T cell exhaustion, leading to ∼40% increase in the exhausted T cell population compared to antigen exposure alone (**Fig. 4D**). Population dynamics of the model suggest a targetable therapeutic window for aKG supplementation, despite the rapid hypoxia-induced exhaustion of Teff cells and insufficiency of immunotherapy alone. Upon integrating these observations, our model predicted that aKG supplementation would augment Teff restoration due to its ability to counteract the detrimental effects of hypoxia (**Fig. 4E-F**). Consistent with our *in vivo* dietary supplementation approach (**Fig. 3F**), just a 1.6-fold increase in serum aKG is predicted to restore effector T cell populations by 5-fold (**Fig. 4E-F**). To apply this prediction to patient data, we assayed serum aKG levels from 10 NSCLC patients, spanning a range from <1 to >25μM (**Supp. Fig. S1N**). Combining our *in silico* model parameterized on experimental data with these clinical aKG levels, we created a metric we termed the “Benej-Mathur index” to identify an NSCLC patient population most likely to benefit from dietary Ca-aKG supplementation based on the serum aKG levels. We used the naturally occurring inflection points in the data to group the patients into low, middle, and high aKG levels (**Supp. Fig. S1N**). Our model predicts that NSCLC patients with serum aKG levels < 8μM will achieve the greatest maximum benefit of Ca-aKG supplementation (**Fig. 4G**).

## DISCUSSION

Loss of effector T cell function and progression of T cell exhaustion are well-established hallmarks of ICI resistance (10, 75). Carrying out the effector function requires a high degree of metabolic plasticity within the nutrient-scarce TIME (76). Recent studies have elegantly demonstrated that TIME-induced metabolic alterations promote free radical-driven impairment of mitochondrial function, leading to suppression of the TCA cycle and loss of the proliferative and cytotoxic capacity (15–17). While this implies that metabolic reprogramming by metabolite supplementation can be a promising therapeutic avenue to restore the Teff function (24), previous studies have shown that clinical translation of metabolite supplementation strategies to improve ICI outcomes is challenging (46).

In the present study, we demonstrate that chronic TcR stimulation decreases the availability of TCA metabolite aKG in a process that is further exacerbated by hypoxia, and that this metabolite ‘starvation’ can be therapeutically exploited to rescue Teff function and feasibly improve ICI response in NSCLC. aKG plays a critical role in many biochemical processes that directly or indirectly affect T cell function including ATP synthesis through TCA cycle, redox metabolism, amino acid synthesis, HIF-1α regulation or its regulatory role in epigenome modulation by affecting H3K27me^3^ demethylation levels (77–80). Our findings support a mechanistic model where exogenous aKG increases the metabolite levels depleted within the TIME, leading to UTX/JMJD3 reactivation and genome-wide methylation decrease evidenced by reduced H3K27me^3^. With no significant changes in superoxide levels, TCA metabolites, amino acid content, or HIF-1α levels in response to aKG supplementation at the concentrations used in our study, the contribution of epigenetic methylation remains a promising mechanism.

A potential limitation is that systemic aKG supplementation is not specific to CD8^+^ T cells only, and will affect other immune cell, stromal and cancer cell components of the TIME. However, immunophenotyping analysis revealed no significant changes in immune cell infiltration of the tumors in either diet cohort, validating that the improved anti-PD-1 response observed in mouse models of NSCLC is not a result of more cytotoxic cells present in the Ca-aKG cohort tumors. Accordingly, *ex vivo* cytotoxicity assay confirmed that antigen-experienced infiltrating CD8^+^ T cells isolated from the Ca-aKG cohort kill cancer cells at a significantly higher capacity compared to CD8^+^ T cells isolated from the regular diet cohort tumors. While this does not necessarily rule out that other immune cell types are not contributing to the synergistic effects of Ca-aKG supplementation on ICI response; combined with *in vitro* data presented herein, these findings confirm that the cytotoxic capacity of CD8^+^ T cells is significantly enhanced in response to increased aKG availability.

This study evaluates Ca-aKG supplementation in the context of cancer immunotherapy in NSCLC. A notable difference in our strategy compared to non-cancer applications of Ca-aKG supplementation is that the synergistic effects of the combination therapy are not dependent upon long-term aKG level elevation, which may not be feasible. The goal of our Ca-aKG supplementation strategy is not to provide anti-tumor effects as a monotherapy. Instead, our approach feasibly targets a transient increase in aKG levels in combination with each dose of immunotherapy to reinvigorate T cell function and help initiate a more robust and sustained immune response.

While we anticipate that many solid cancers would benefit from this dietary supplementation strategy, our findings imply that NSCLC patients may be particularly responsive for multiple reasons: 1) serum aKG levels were shown to naturally decline with increasing age (70); 2) NSCLC disproportionately affects older individuals (median age is 71 years); 3) serum aKG levels decline in older NSCLC patients; and 4) reduced serum aKG levels correlate with poor overall survival in NSCLC patients treated with ICIs. These findings also challenge the current standard of using young, typically 7-week-old mice for preclinical research in NSCLC, as the serum aKG levels in young mice are significantly higher compared to aged mice, and as such, given the role of aKG in ICI response, do not reliably model the TIME conditions typically occurring in NSCLC patients. Instead, we propose to adopt aged mouse models as the “gold standard” for NSCLC immunotherapy research.

Dietary aKG supplementation in non-cancer healthcare applications has been found to be well-tolerated; typically, only mild gastrointestinal side effects such as bloating and diarrhea were reported upon long-term use (81). In this study, we used lung adenocarcinoma-derived CMT167 cell line that harbors an activating *Kras* mutation and is *p53/Stk11* wild-type; and squamous cell carcinoma-derived KLN 205 cells that are *Kras/Stk11* wild-type and *p53* mutant (82, 83). Our initial findings validate that cell-permeable aKG supplementation alone does not affect the proliferation of CMT167 or KLN 205 cells *in vitro*; and the 4% Ca-aKG-enriched diet does not increase tumor growth *in vivo*, as documented in both male and female, young and aged mouse hosts bearing CMT167 or KLN 205 tumors. While this suggests that the histological subtype, genomic background, sex or age should not be an exclusion factor for a potential Ca-aKG supplementation strategy, further studies will be needed to evaluate the role of these factors in modifying the aKG levels within the TIME and assess possible response to Ca-aKG supplementation. To further support the safety of this approach, initial complete necropsy and blood analysis showed no toxicities in mice on 4% Ca-aKG diet compared to regular diet. While our findings support the feasibility of the dietary supplementation-based augmentation of ICI response, future studies are needed to fully evaluate the dose response and establish the dosing associated with maximum effect on ICI response.

In conclusion, preclinical and clinical findings presented warrant further investigation to support translation into the clinic, especially since Ca-aKG and other cell-permeable aKG variants are already widely available without a prescription and patients may already be taking them, making it all the more important to fully investigate the mechanism and lay the groundwork for future studies.

## METHODS

### Cell Lines

All cell lines were purchased from the American Type Culture Collection (ATCC) and grown in DMEM (Gibco) supplemented with 10% FBS (Gibco) and 1% Penicillin-Streptomycin (Gibco). Cell counts and cellular proliferation were established using automated cell counter to determine the cell number and trypan blue exclusion assay to establish the fraction of viable/dead cells. All cell lines were authenticated using short tandem repeat (STR) profiling within the past 3 years. All experiments were performed with mycoplasma-free cells.

### Animal experiments

The CMT167 cells (1.25 × 10^6^) were injected s.c. into the flanks of 7-week-old or 35-week-old female or male C57BL/6J mice; or the KLN 205 cells (2 × 10^6^) were injected into the flanks of 7-week-old or 35-week-old female or male Dba/2 mice following Institutional Animal Care and Use Committee (IACUC)-approved protocols. Caliper measurements of opposing diameter were used to calculate the tumor volumes. For the Ca-aKG dietary supplementation, 4% Ca-aKG diet was formulated in regular Teklad 7912 diet and irradiated at Inotiv Laboratories. Unless stated otherwise, mice were randomized into groups fed with regular diet or Ca-aKG diet when tumors became palpable and were maintained on the respective diets for the entire duration of the experiments. For the immunotherapy response evaluation, upon reaching 150 mm^3^, the mice received three doses of anti-PD-1 antibody (70 μg, i.p. against rat IgG isotype control, BioXcell). Tumor volumes were measured until the posttreatment volume increased threefold.

### *In vitro* T cell exhaustion assay

Primary naïve CD8^+^ T cells were extracted from mouse spleens using EasySep Mouse CD8a Positive Selection Kit II (STEMCELL Technologies #18953). Naïve CD8^+^ T cells were activated for 72 hours using anti-CD3/CD28 monoclonal antibodies (Invitrogen #16-0032-86 and #16-0281-86, respectively) in RPMI 1640 medium (Gibco) containing 25 mM HEPES (Gibco); 1x MEM Non-Essential Amino Acids (Gibco); 50 μM 2-mercaptoethanol (Millipore Sigma) and 20 ng/ml interleukin-2 (Millipore Sigma). Rapid *in vitro* exhaustion using chronic TcR stimulation with 3 μg/ml anti-CD3 antibody was adapted from (53). Briefly, following activation, T cells were re-stimulated with fresh anti-CD3 antibody every two days under normoxic or hypoxic conditions (1% O_2_) ± 10 mM dimethyl alpha-ketoglutarate (Millipore Sigma). On Day 10, the T cells were harvested for downstream analyses. These include: multispectral flow cytometry analysis by Cytek Aurora instrument using our validated panel of 32 lineage and effector/exhaustion markers (CD45, CD3, CD8, CD4, FoxP3, CD11b, PD-1, Tim3, Slamf6, TOX, Tcf-1, CD44, CD62L, CTLA-4, Lag-3, Klrg1, T-bet, Ki-67, Bcl2, EOMES, Vista, TIGIT, CX3CR1, ICOS, CD27, CD38, CD95, NK1.1, CD25, GITR, CD69, GzmB); flow cytometry using anti-TOX-PE (Miltenyi Biotec) and anti-TCF1/7-Alexa Fluor 647 (Cell Signaling); and bulk RNA-Seq analysis by a commercial end-to-end service.

### Mitochondrial fuel utilization assay

Activated CD8^+^ T cells were incubated for 10 days under chronic TcR stimulation in normoxic or hypoxic conditions. On Day 10, the cells were plated into a MitoPlate S-1 Assay (Biolog, #14105) plate at 35,000 cells/well. Colorimetric changes representative of electron transport chain-mediated reduction of a tetrazolium redox dye were imaged after three hours following manufacturer’s instructions.

### Quantification of effector cytokine production

Activated CD8^+^ T cells were incubated for 10 days under chronic TcR stimulation in normoxic or hypoxic conditions ± 10 mM cell-permeable aKG. On Day 10, the cell culturing media were collected and extracellular TNFα and IFNγ levels were assayed using Mouse TNF-alpha Quantikine ELISA Kit (R&D Systems, #MTA00B) and Mouse IFN-gamma Quantikine ELISA Kit (R&D Systems, #MIF00), respectively, following manufacturer’s instructions. Absolute extracellular effector cytokine quantities were normalized to endpoint cell count for each condition.

### UTX/JMJD3 activity assay

Activated CD8^+^ T cells were incubated for 10 days under chronic TcR stimulation in hypoxic conditions ± 10 mM cell-permeable aKG. On Day 10, the cells were harvested and 2×10^6^ were assayed for nuclear extraction using Nuclear Extraction Kit (Abcam, #ab113474) following manufacturer’s instructions. Obtained nuclear extracts were quantified using Pierce BCA Protein Assay (Thermo) and 2.5 μg of each sample was assayed in triplicates to determine the ability of UTX/JMJD3 to remove plate-bound H3K27me^3^ using *kdm6a/kdm6b* Activity Quantification Assay Kit (Abcam, #ab156910) following manufacturer’s instructions.

### Quantification of metabolite levels in biological samples

The levels of aKG in murine T cells and human NSCLC patient serum samples were identified using commercial colorimetric assay (α-Ketoglutarate Detection Assay Kit, Cayman #701350) following manufacturer’s instructions. Prior to the assay, serum samples were centrifuged through 10 kDa molecular weight cut-off filter spin columns (Abcam #ab93349). Additional mouse serum, human healthy blood donor serum samples and archival NSCLC patient serum samples were analyzed using liquid chromatography-mass spectrometry (LC-MS) at the Campus Chemical Instrument Center’s (CCIC) Mass Spectrometry and Proteomics Facility. For mass spectrometry imaging (MSI), an HTX M5 sprayer was used to deposit a layer of MALDI matrix to optimize for TCA cycle metabolites in negative mode. Pixel-by-pixel MS intensity was analyzed using Bruker analysis tools to produce spatially resolved chemical images of tumor tissues.

### Quantification of extracellular lactate levels

Activated CD8^+^ T cells were incubated for 10 days under chronic TcR stimulation in hypoxic conditions ± 10 mM cell-permeable aKG. On Day 10, the cell culturing media were collected and extracellular lactate levels were assayed using L-Lactate Assay Kit (Millipore Sigma, #MAK329) following manufacturer’s instructions. Absolute extracellular lactate quantities were normalized to endpoint cell count for each condition.

### *Ex vivo* cytotoxicity assay

To determine the cytotoxic activity of infiltrating T cells, CMT167 cells (1.25 × 10^6^) were injected s.c. into the flanks of 35-week-old female or male C57BL/6J mice; or KLN 205 cells (2 × 10^6^) were injected into the flanks of 35-week-old female or male Dba/2 mice following Institutional Animal Care and Use Committee (IACUC)-approved protocols. When tumors became palpable, mice were randomized into groups fed with regular diet or 4% Ca-aKG diet. After two weeks, tumors were harvested, homogenized, and infiltrating CD8^+^ T cells were extracted using EasySep Mouse CD8a Positive Selection Kit II (STEMCELL Technologies #18953). Spleens from tumor-bearing mice were used to extract naïve CD8^+^ T cells as a control. Dead T cells were removed using the EasySep Dead Cell Removal AnnV Kit (STEMCELL Technologies #17899). One day prior to infiltrating T cell extraction, CMT167 or KLN 205 cells were plated at 15,000 cells/well into Agilent xCELLigence RTCA S16 platform and allowed to reach exponential phase. After reaching exponential phase, extracted antigen-experienced infiltrating CD8^+^ T cells were added to the proliferating CMT167 or KLN 205 cells at a 3:1 effector-to-target ratio. The cytotoxic capacity of extracted infiltrating T cells was evaluated as a function of CMT167 or KLN 205 proliferation. Obtained Cell index values were validated using trypan blue exclusion assay confirming observed cancer cell kill rates.

### Single-cell western (scWestern) analysis

Activated CD8^+^ T cells were incubated for 10 days under chronic TcR stimulation in hypoxic conditions ± 10 mM cell-permeable aKG. On Day 10, T cells were harvested and pooled onto an scWestern chip at a concentration of 1 x 10^5^ cells/ml. On-chip chromatin digestion consisted of permeabilization in 0.025% L-α-lysophosphatidylcholine (LPC) in permeablization solution #1 (150 mM sucrose, 80 mM KCl, 35 mM HEPES, 5 mM K_2_HPO_4_, 5 mM MgCl_2_, 0.5 mM CaCl_2_, pH 7) at 37 °C for 2 minutes; followed by a wash in permeablization solution #1 (without LPC) and incubation with 300 U MNase + 250 U DNase in permeabilization solution #2 (150 mM sucrose, 50 mM NaCl, 50 mM Tris-HCl, 2 mM CaCl_2_, pH 7) at room temperature for 5 minutes. Histone H3 and H3K27me^3^ levels were determined by scWestern analysis using ProteinSimple Milo Single-Cell Western Platform (Bio-Techne) using validated antibodies (Cell signaling #4499 and Abcam #6002, respectively), paired with Alexa Fluor 647- (for Histone H3 Invitrogen #A27040) or Alexa Fluor 555 (for H3K27me^3^ Invitrogen #A31570)-conjugated secondary antibodies. scWestern images were obtained using Innoscan 710 Microarray Scanner (Innopsys) and analyzed by Scout Software Package (ProteinSimple).

### Human samples

Samples were obtained from two cohorts. The first cohort was NCT06630429, which is a prospective, observational study of patients undergoing anti-cancer immunotherapy. Eligible patients were receiving immunotherapy as part of standard care and had no contraindications to phlebotomy. After consent and eligibility confirmation, patients underwent serial blood sample collection at each of the first 4 infusion visits and at 3, 6, 12 and 24 months.

The second cohort is the retrospective Total Cancer Care (TCC) protocol. TCC establishes a unique collection of blood, tissue, other biological samples and associated data (e.g., medical records data) from thousands of patients with cancer or at risk of having cancer. Through the consent, patients provide permission to study their medical and related records, to collect: excess tissues or body fluids removed at the time of planned surgery or biopsy, tissue previously collected from the patient that is being stored (if available), blood in conjunction with planned clinical blood draws or drawn specifically for this study, and other non-invasive specimens. Tissues (normal and tumor) or fluids may be collected at the time of standard of care surgical procedures and biopsies or autopsy if the autopsy consent form has been signed for future cancer research studies.

### Model Development

We model a population which has *N* cells of three kinds: effector T cells (effT), progenitor exhausted T cells (proT) and terminally exhausted T cells (exT). Population *N* has three spatially segregated sub-populations, *n_1_*, *n_2_*, and *n_3_*, each containing effT, proT, and exT cells:

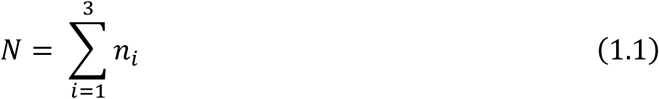

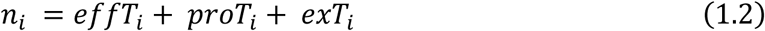

We constrain the model to a fixed number of cells, such that *N* and *n_i_* are constants. This imposes the assumption that there is no growth nor death of T cells, nor influx/efflux into the system(s). However, we allow transitions between the cell types as follows:

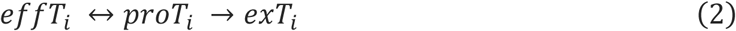

If *n_i_*[*t*] is the number of cells in a given population at time *t*, we can describe each subpopulation with the following system of ordinary differential equations:

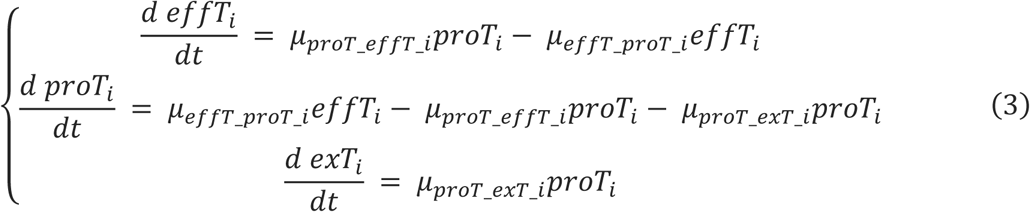

Where *μ_a_*__*b*_*i*_is the transition rate from cell type *a* to cell type *b* in *n_i_*.

### Model Application

We allow nonnegative values of *μ_a_*__*b*_*i*_. To parameterize starting conditions, we impose *N =* 500, representing approximately 5mm^2^ of a T-cell dense lung tumor (84). We consider the 3 subpopulations to be distinct regions of varying hypoxia. Buffa hypoxia scores for NSCLC patients can be partitioned into values of -3 (quartile 1), 19 (quartile 2), and 31 (quartile 3), representing increasing levels of hypoxia. Given a slight right skew and positive median value, we assume *n_1_*= 100, *n_2_* = 200, *n_3_* = 200 for quartiles 1 to 3, respectively. This yields simulated T cells in a tumor that leans hypoxic.

#### Modeling the Effect of Hypoxia on T cell dynamics

We consider antigen exposure and hypoxia as variables that drive the forward reactions of T cell exhaustion, and at baseline assume equal effect of the variables on *μ_effT_*__*proT*_*i*_ and *μ_proT_*__*exT*_*i*_. For all *n_i_* we assume the effect of antigen alone as *μ_effT_*__*proT*_*i*_ = *μ_proT_*__*exT*_*i*_ = .05 . Given that hypoxia must reach a certain threshold to have an effect, we scale the hypoxia quartiles to affect transition rates as follows:

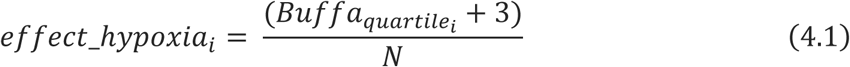

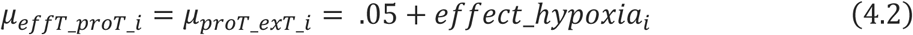

At *t* = 0 we assume all cells are effT.

#### Modeling the Effect of aKG on T cell dynamics

We consider immunotherapy and aKG supplementation as variables that drive the reverse reaction of T cell exhaustion. We assume immunotherapy leads to a *μ_proT_*__*effT*_*i*_ that counteracts the antigen contribution to *μ_effT_*__*proT*_*i*_. We assume aKG supplementation reduces *μ_effT_*__*proT*_*i*_ proportional to its value. We parameterize this using data from Figure 3, and consider supplementation to increase aKG levels from *jj* = 1X to 2X fold change (*aKG_FC_*) in intervals of 0.1. We assume a nonlinear relationship between aKG levels and its effect on T cell transitions as follows:

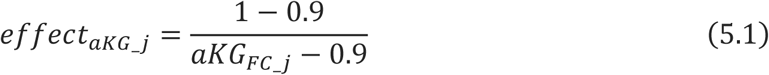

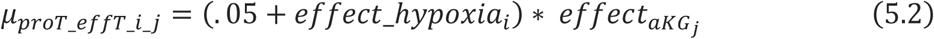

Such that *effect_aKG_*__*jj*_ asymptotically reaches .9 (**Fig. 1E**) and affects each *n_i_* equally. To model the start of immunotherapy and aKG supplementation after T cells have started to become exhausted, we assume that at *t* = 0, the *n_i_* population distribution is that of *t* = 25 (arbitrary units) in the absence of treatment, during the window where progenitor T cells are linearly increasing.

#### Predicting aKG Supplementation Effect on Patient’s T cell populations

We observe 2 inflection points in patient aKG level data. The mathematical inflection point, when *f*^′′^ = 0, and the visual inflection point when aKG rapidly increases, occur at aKG values of ∼ 8 and 12, respectively. We therefore partitioned patients into 3 groups: aKG<8, 8<aKG<12, and aKG>12. We then extrapolated aKG supplementation levels from *jj* = 1X to 3X to capture this range, and calculated the relative effect on predicted effector T cell restoration in each patient group.

## ACKNOWLEDGEMENTS

This work was supported by funding from National Institutes of Health / National Cancer Institute grant K22 CA282363 (MB) and OSUCCC Startup Fund (MB). The authors would like to acknowledge assistance from the OSUCCC Clinical Translation Science Shared Resource, the OSUCCC Hematology Tissue Bank Shared Resource, the OSUCCC Flow Cytometry Shared Resource, and the Campus Chemical Instrument Center.

Research reported in this publication was also supported by the OSUCCC and the National Institutes of Health under grant number P30 CA016058. We thank the Total Cancer Care of the Biospecimen Services Shared Resource at The Ohio State University Comprehensive Cancer Center, Columbus, OH, for specimens and data. The results shown here are in part based upon data generated by the TCGA Research Network: https://www.cancer.gov/tcga.

NCT06630429 is supported by NIH/NCI (R01CA273924) and the LUNGevity Foundation. Figures were created with <u>BioRender.com</u>

