## Supplementary Data for "Targeting alpha-ketoglutarate starvation within the tumor-immune microenvironment rescues T cell effector function in cancer immunotherapy"

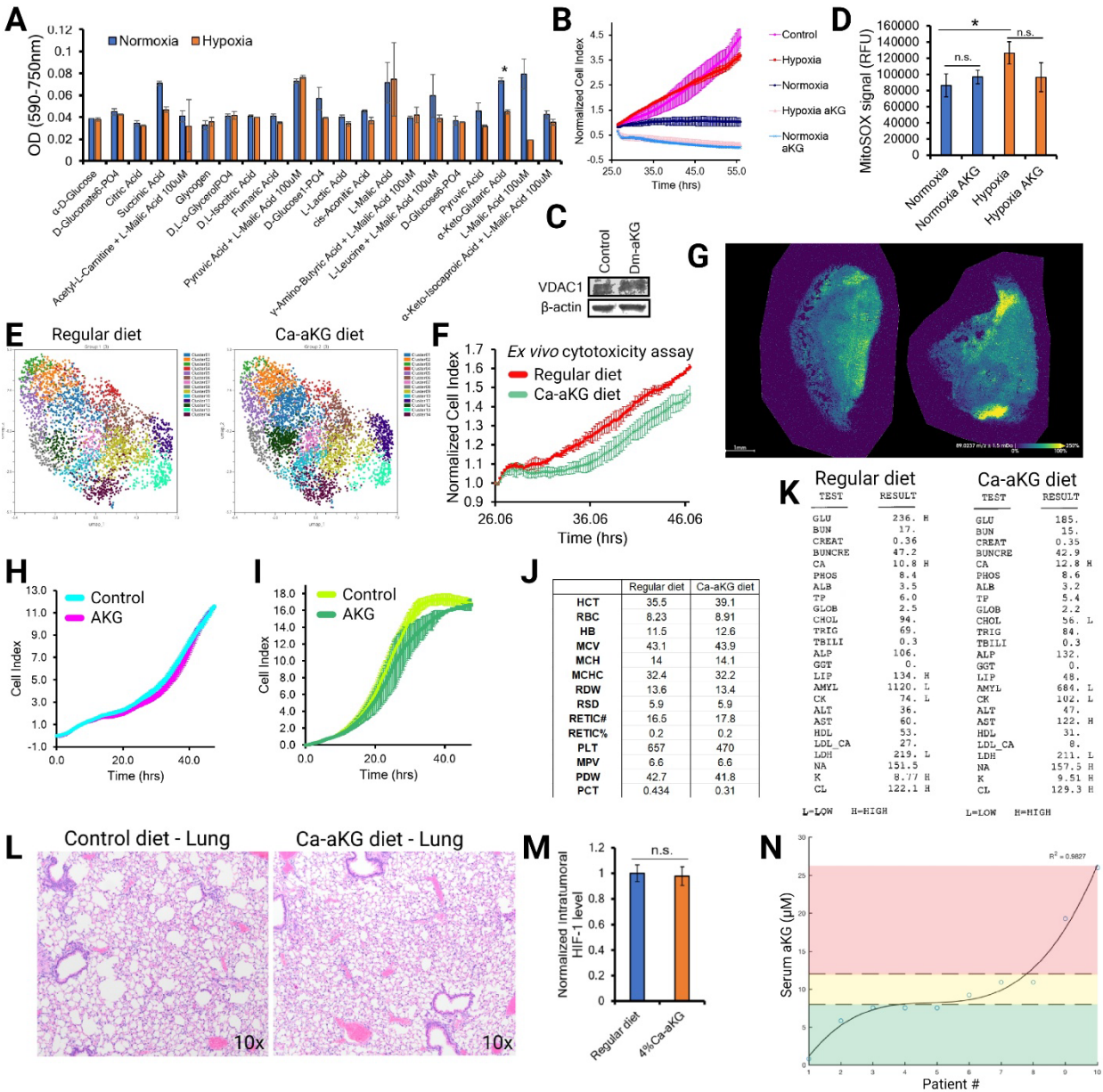

Regular diet (*red*) or Ca-aKG diet (*green*) cohort KLN 205 tumors grown in Dba/2 mice. Curves represent KLN 205 proliferation when co-cultivated with antigen-experienced T cells in a 3:1 effector-to-target ratio, as determined by Agilent xCELLigence RTCA S16 platform (n=5). **(G)** Representative MALDI-ToF-based spatial metabolomics analysis of CMT167 tumor sections from mice fed with regular diet (*left*) or 4% Ca-aKG diet (*right*). Signal intensity depicts mass spectrum distribution of lactate. **(H-I)** Proliferation of CMT167 (H) or KLN 205 (I) cells ± 10 mM cell-permeable aKG, as determined by Agilent xCELLigence platform (n=3). **(J-K)** Representative blood analysis results from aged mice fed with regular diet or 4% Ca-aKG diet for 2 weeks. (n=4-5) **(L)** Representative H&E staining of lung tissue from CMT167 flank tumor-bearing C57BL/6J mice fed with regular diet or 4% Ca-aKG diet for 2 weeks. **(M)** Intratumoral HIF-1α levels in CMT167 tumors harvested from aged mice fed with regular diet or 4% Ca-aKG diet for 2 weeks, as determined by IHC staining. **(N)** Inflection points in the NSCLC serum aKG data used to group the patient samples into low, middle, and high aKG levels.
